# Carrying oversized loads may create pheromone “blind spots” in leafcutter ants

**DOI:** 10.1101/2024.12.11.628008

**Authors:** Katherine Porras-Brenes, Sabrina Amador-Vargas

## Abstract

Understanding animals’ sensorial abilities and cognitive processes while foraging helps explain why animals depart from theoretical optimal foraging. Here, we studied foraging decisions in leafcutter ants (*Atta colombica*), which form and follow pheromone foraging trails that workers smell by tapping the substrate with the antennae. Carrying oversized items transports more plant material in a single trip, but workers walk more slowly and can delay the nestmates walking behind them. We tested the hypothesis that balancing an oversized load limits the ability to tap the ground with the antennae, therefore reducing the ability to smell the foraging trail. As expected, we found that the number of antennae taps per step was (1) fewer in laden vs. unladen workers, (2) fewer as loads increased in area, but only for larger ants, and (3) unrelated to the load shape. Second, workers increased the antennae taps and speed after experimentally reducing standardized loads. Last, we evaluated the allometric relation between the antennae length and worker size, and found that it showed negative allometry. Hence, larger ants had proportionally shorter antennae, which could explain why larger workers are more impacted by oversized loads in the number of antennae taps. Overall, our results support that carrying an oversized load limits the ability of workers to tap the ground, analogous to the blind spots in a large truck. Our findings reveal a sensory limitation to foraging and reflect how foraging decisions in a social organism are adjusted to the individual capacities and the colony needs.

## INTRODUCTION

Foraging is a behavior essential for the individual’s fitness and relies on a set of skills that vary depending on the individual’s abilities and on the environmental limitations to develop those behaviors. Foraging decisions are made based on the information that the animal’s sensory system gathers (Hemingway et al., 2018). Also, foraging decisions can also be limited by morphological or physiological constraints, independent of their sensory system (e.g., Friedlaender et al., 2016). Hence, understanding the sensorial abilities and cognitive processes of animals helps explain how animals decide and why they depart from theoretical optimal foraging.

Living in a group could also influence foraging decisions. Foraging in a social context requires that decisions balance the individual and the group’s needs. Foraging in a collaborative society implies communication, as well as organization, and sometimes memorization (Dukas & Real, 1991). Among social animals, eusocial insects are an outstanding example of social foraging. Foraging in eusocial insects has two key differences from solitary organisms with implications for their sensory systems. First, social insects mostly collect food for the brood instead of only fulfilling their needs (Brian, 2012). This creates the need to balance between foraging decisions that benefit the individual and decisions that benefit the entire group. Second, eusocial insects are central place foragers, i.e., they bring food to a particular location, which requires navigating back to the nest (Bell, 1990; Narendra, 2021). For both reasons, the foraging effort relies on communication among nestmates. Hence, in social insects, foraging behavior includes sensing the environment and nestmate cues, and it also requires that the foraging decisions do not interfere with their ability to navigate back to the central place.

In the neotropics, leafcutter ants perform a massive foraging operation that constitutes an excellent subject to understand foraging decisions in a eusocial organism. Workers are organized by castes with a set of specific functions, and forage for plant material to feed a fungus garden (Hölldobler & Wilson, 2010; Calheiros et al., 2019; Hubbell et al., 1980). A key foraging decision that leafcutter ant workers make is regarding load size. Carrying the maximum load according to their muscular force has advantages and disadvantages for the colony and the individual. Accordingly, workers often carry less than the maximum weight they can lift (Segre & Taylor, 2019). Load size decision can be determined by environmental factors (e.g., wind: Alma, Farji-Brener & Elizalde, 2016; and rain: Farji-Brener et al., 2018), social conditions (e.g., flux density in the trail: Farji-Brener et al., 2011; or information transfer needs: Roces & Bollazzi, 2009), or internal conditions (e.g., motivation and energy: Püffel et al., 2021; Roces, 1993; Roces, 2002). Carrying a large load would optimize the amount of material brought in a single foraging trip. However, depending on the conditions, bringing a large load can slow the workers’ pace and even impact the speed of workers walking behind, thus affecting the entrance rate of material to the nest (Rodriguez-Planes & Farji-Brener 2019). Farji-Brener et al. (2011) found that workers carrying oversized loads (called “truck drivers”) are scarce when trail traffic is high. Hence, a balance is achieved between the individual benefit and a reduced cost for the entire colony. Carrying oversized loads can be mechanistically challenging for the worker, even though larger loads optimize the material brought to the nest in one trip. Here, we considered yet another possibility that explains the load size decision: how the load may interfere with the organism’s sensory system.

In our study, we tested the hypothesis that carrying a load negatively affects the capacity of leafcutter ants to tap the ground with the antennae to detect the foraging trail. Leafcutter ants in the genus *Atta* form well-defined foraging trails that are maintained by laying pheromones on the trail surface (Guerrero, 1988). Nestmates detect the chemical signals using their antennae and lay more pheromones, reinforcing the chemical trail (Evison, Hart & Jackson, 2008; Kleineidam et al., 2007). Those pheromones help the ants navigate from the nest to the plant source and back (Hölldobler & Wilson, 1990; Hart & Jackson, 2006; Evison et al., 2008), and require the ants to tap the ground with the antennae to smell the pheromones. While carrying a large load, maintaining balance may cause the ant to keep the head in a position that impedes constant tapping of the ground with the antennae. Thus, a large load may hamper the perception of chemical signals and may encourage the worker to find another way to orient back to the nest. The situation is analogous to the blind spots of large trucks moving goods. Trucks transport a large mass in a single trip, but the load interferes with the driver’s visibility (Umar et al., 2022), and the truck moves forward despite being unable to see all the surroundings. Oversized loads in ant workers could also cause “pheromone blind spots”, if they limit the worker’s ability to tap the ground with the antennae.

As it is known that unladen ants interact with laden ants in the trails and use that information to search for specific foraging items (Burd & Aranwela, 2003; Dussutour et al., 2007; Farji-Brener et al., 2010). Hence, we also evaluated whether ants carrying oversized loads were tapping other nestmates with the antennae, rather than the ground, to keep on the trail. As other factors can influence the load size in field conditions, we also experimentally offered standardized foraging loads to the ants that were cut while the ants were walking, to assess the effect of the load reduction on the number of antennae taps. Finally, we studied whether the relation between antennae and body size in leafcutter ant workers is allometric, as this could determine whether ants of all sizes have similar limitations to lower the antennae while carrying a large load.

## METHODS

### Study site and study species

We conducted our study in Gamboa, Colón, Republic of Panama (9° 7’ 30” N, 79° 42’ 00” W) from May to July 2022. The area is a tropical humid forest, 30 m.a.s.l., in the Panama Canal watershed (Serrano, Nuñez & Valleter, 2017). Rainfall is typically 2000 and 3000 mm per year, and temperatures vary between 24 and 27°C (Valderrama, Herrera & Salazar, 2008). We worked in 20 mature nests of the leaf-cutter ants *Atta colombica.* Mature colonies of these ants have millions of polymorphic workers, which forage on a foraging trail system with well-defined main trails and secondary branches (Wirth et al., 2003). In general, we explored three lines of evidence: (1) we evaluated the sensorial capacity of the ants according to the load by counting the antennae taps per step, and relating them with load area and shape in field foraging ants. We also explored tapping nestmates as a potential information source in these natural trails. (2) We performed an experiment in the field, in which we reduced the load to test if the antennae taps per step decreased after the reduction. Finally, (3) we explored the allometric relation between antennae length and worker size. All the statistical analyses and graphs were done using R version 4.3.1 (R Core Team, 2017). Videos were recorded using a Digital Sony 4k Video Camera Recorder FDR-AX53.

### Sensorial capacity according to the load

In each nest, we recorded workers on main trails that facilitated recording at the level of workers (e.g., sidewalks and roads). The ants were recorded with the camera placed 5 cm from the trail. We recorded trails with low or moderate worker flux during the day, as they have higher chances of finding workers with oversized loads (Farji-Brener et al., 2011; Pereyra & Farji-Brener, 2020).

We video-recorded workers carrying different load sizes from the side at ground level, trying to focus their legs and antennae. After they passed the video shooting frame, we collected the loads to photograph and measure them. We measured the workers’ head width and released them back on the trail. We only collected data for workers carrying leaf fragments and excluded workers carrying other materials (e.g., fruits and flowers). We also recorded unladen workers walking outbound (away from the nest).

#### Antennae taps and load presence

To test the hypothesis that the presence of the load negatively influences the ability of workers to tap the trail with the antennae (hence, detecting the chemical trail), in the videos we counted the number of times the antennae tapped the ground and the number of steps that the ants did on five centimeters of trail. We related the number of antennae taps (response variable) with the presence of the load (fixed factor) using a Generalized Linear Model (GLM) with Poisson distribution, and including the number of steps as a covariate or random factor. In addition, a chi-squared test was performed to evaluate whether the absence of antennae tapping was more frequent in laden ants.

#### Antennae taps per step and load size and shape

First, we characterized the load shape as more rounded or more elongated, according to how the ant worker carried it. Load length was measured upwards at the maximum distance from the point in the leaf fragment where the ant was holding it with the mandibles (Fig S1). Then load width was calculated as the longest line perpendicular to the length. Then, we combined both variables as load shape using a width-to-length index (WL index), in which values close to one indicate rounded loads, values above one indicate wider loads, and values below one indicate more elongated loads (Shi et al., 2021). Second, we calculated the leaf area in Image J (Abràmoff et al., 2004) using the calibrated picture. The load size and shape measurements were blind to the size of the ant, or the number of antennae taps per step.

As the number of steps increased with the number of antennae taps (GLM: Estimate=0.081, z=19.01, p<0.0001), we used the number of antennae taps per step as our response variable for the following analyses. Since all workers have different sizes and walk at different speeds, we considered that the number of steps before making another tap with the antennae is the best estimate of how much a worker can move forward before smelling the trail again. We used a GLM to evaluate whether the number of antennae taps per step (response variable), was influenced by an interaction of load size (fixed factor) and worker size (ant head width; fixed factor). Further, to evaluate the effect of load shape in the antennae taps per step, we ran a separate GLM with the interaction between the leaf shape, i.e., the WL index (fixed factor) and the workers’ head width (fixed factor) on the number of antennae taps per step (response variable). Finally, we used data from unladen ants to evaluate whether the antennae taps per step (response variable) are related to worker size (fixed factor) using a GLM with a Gaussian distribution.

#### Antennae taps on nestmates

As workers may detect the foraging trail by tapping the nestmates with the antennae rather than the ground, we counted the number of nestmate interactions between laden and unladen ants. We predicted that ants with relatively larger loads would contact the nestmates more often than workers carrying relatively smaller loads. A nestmate interaction was defined as the contact between the antennae of the laden worker and any part of a nestmate. We then used a GLM with Poisson distribution using the number of nestmate interactions as the response variable, and the interaction between load area and head width (fixed factors).

### Load reduction experiment

To experimentally determine whether load size is determining the number of antennae taps on the ground, we conducted an experiment where the same ant was measured before and after reducing the load size. First, we offered loads of known sizes made of cellulose filter paper (Whatman 2.3 cm diameter), which were dipped in orange juice to make them more attractive to ants (Moll, Roces & Federle, 2013). After they were dry, we cut them in half across the diameter and used a pencil to mark a midline to cut them later through the radius during the experiment. We offer the prepared papers as a pile next to a foraging trail. Once a worker picked up the filter paper, we video recorded it during 5 cm of the trail. Then, we reduced the load size to half by carefully cutting it with scissors while the ant was carrying it. We let the ant continue walking and recorded the same worker farther in the trail with the reduced load for 5 cm (Fig. 1). The area of the initial offered load (before cutting) was 2.07 cm^2^, and after reducing the size, the loads were 1.04 cm^2^. Compared with the results from the previous observations, the loads offered initially were similar to an oversized load in nature, whereas reducing the experimental loads by half, made them fall in a medium-sized load in nature (Fig. S2). In other words, at first, the ants were “truck drivers,” and after cutting the fragment, they were not (Farji-Brener et al 2011). As a control, we did a similar experiment, but we covered the sharp side of the scissors with masking tape. We then touched the experimental load with the scissors (which were unable to cut) and recorded the ants before and after the manipulation.

**Figure 1.**
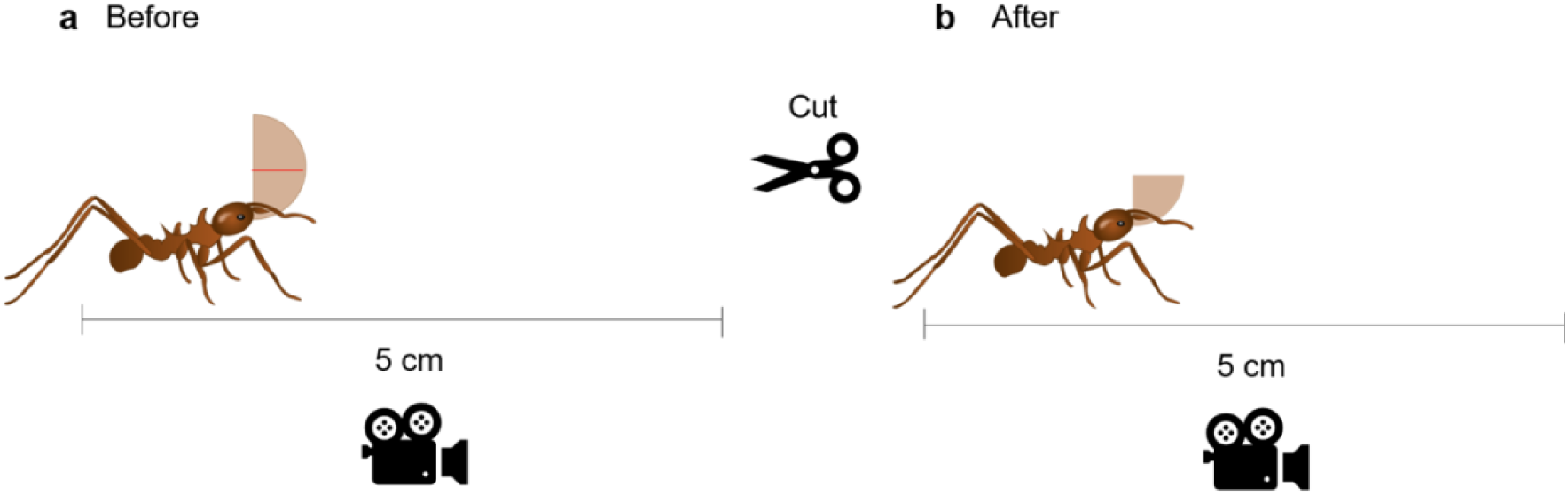
Experimental reduction of load size in leaf cutter ants (*Atta colombica*). **(a)** Before reducing the load size, workers lifted a 2.3 cm diameter Whatman paper piece, and were recorded while walking 5 cm of the trail. **(b)** Then, the load was reduced in half, and the worker was recorded while walking 5 cm of the trail. We compared the number of antennae taps on the ground per step and the walking speed at the two load sizes.

From the videos, the number of steps and the number of antennae taps were counted before and after manipulation for each worker. As the manipulation of the load is evident in the video, these counts could not be made blind to the treatment. We also calculated the walking speed using the time the ant took to walk the 5 cm of trail visible in the videos. We analyzed this data using a Generalized Linear Mixed Model (GLMM) to assess the effect of the treatment (before and after reducing load size, or before and after touching the load with the dull scissors) on the number of antennae taps per step (response variable), and including the individual worker as the random factor (the paired observations), and the workers’ head width as a covariate. We performed a similar analysis but using ant walking speed as the response variable.

### Allometry of antennae length

We also explored the allometric relation between antennae length and worker size (head width or thorax length), because this may hint at whether workers with proportionally longer antennae, would be able to carry proportionally larger loads, as they presumably could tap on the ground while not lowering the head as much as they would with shorter antennae. Hence, we measured ant worker head width, thorax length, and antennae length using calibrated photographs taken in a stereoscope, using Image J (Abràmoff et al., 2004). Body parts measurements were done by a person unaware to the tested hypothesis (blind recording of observations). We used Ordinary Least Square regression in the “sma” R package (Warton et al., 2012) to calculate whether the slope of the regression between the log-transformed measure of the body segments (antennae or leg length) and the body size (thorax length and head width) was different from 1. In the case of antennae, negative allometry (slope < 1) would indicate that larger ants have proportionally shorter antennae than smaller ants. Positive allometry (slope > 1) would indicate that larger ants have proportionally longer antennae, and a slope = 1 would indicate isometry. As we wanted to make sure that any allometric relation was specific to the antennae, we also measured the length of the first and third leg (total length of tibia, femur, and tarsus), and calculated the allometric relation of those structures as well.

### Ethical note

Ethical Note: The study complies with the laws of the Republic of Panama under MiAmbiente permit ARB-128-2022. All our observations occurred in the field, and the ant workers returned to their colonies after field measurements. Ants for the allometry data were cold euthanized before storing in 70% ethanol.

## RESULTS

### Sensorial capacity according to the load

#### Antennae taps and load presence

We observed 813 laden ants and 799 non-laden ants from 20 nests. On average, loaded ants performed fewer antennae taps (2.40 ±1.75), than unladen ants (4.72 ±2.06; Estimate=-0.77, Z _(1611)_=-27.21, *P* <0.0001; Fig. 2), while controlling for the number of steps which are positively correlated with the number of antennae taps (Estimate=0.081, Z=19.01, *P* <0.0001). Moreover, laden ants more often walked the studied distance without making any antennae tap, compared with unladen ants (X^2^=85.53, d.f.=1, *P* <0.0001).

**Figure 2.**
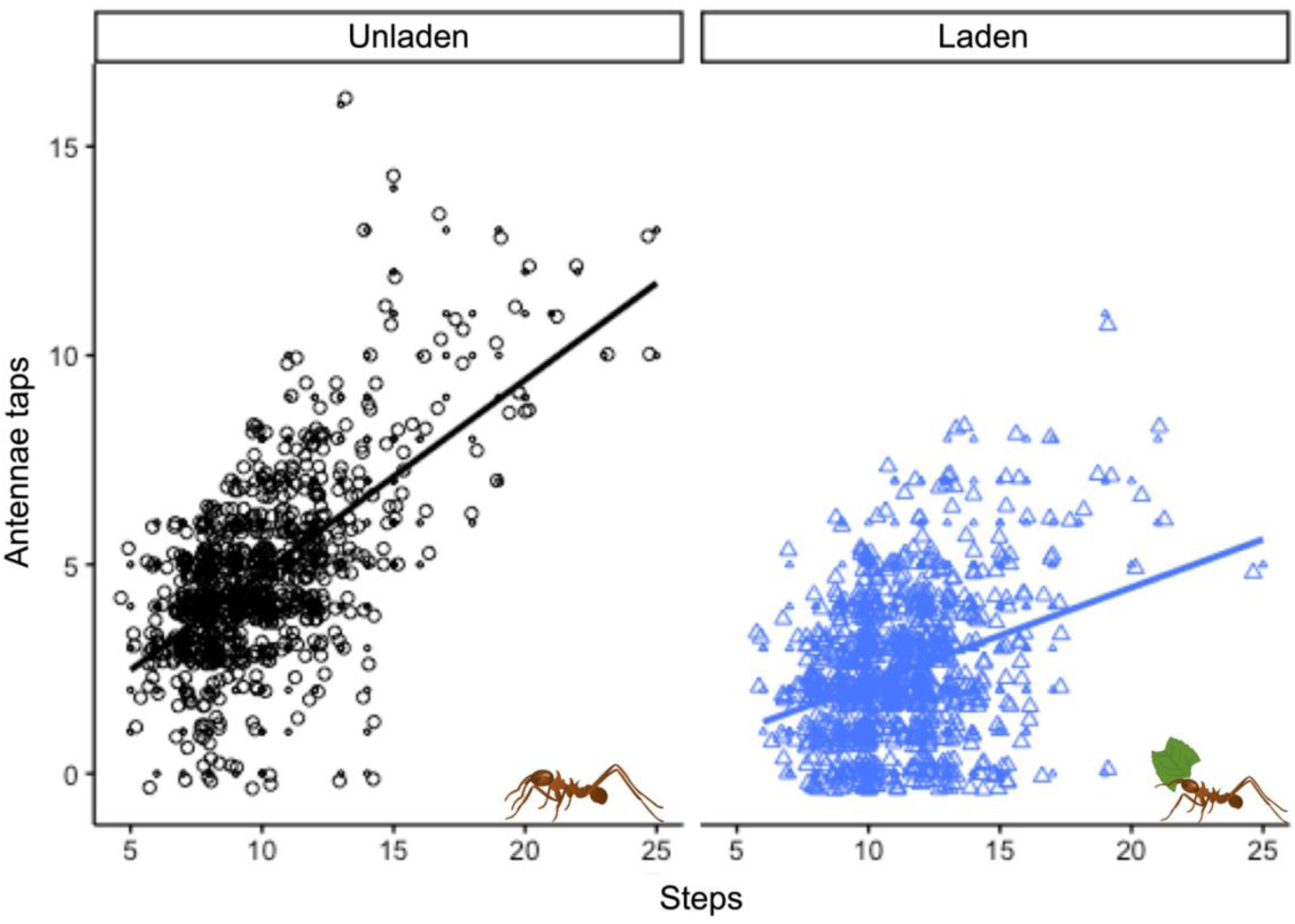
The number of antennae taps in leafcutter ants (*Atta colombica*) increased with the number of steps in (a) unladen workers, and (b) this relation is affected by carrying a load. Symbol size relates to the number of observations with the same value.

#### Antennae taps per step and load size and shape

The number of antennae taps per step was affected by an interaction between load area and head width (Estimate = -0.63, t_(808)_ = -1.90, *P* =0.05). This means that the effect of the load area on the number of antennae taps depends on the ant size: larger but not smaller ants have fewer antennae taps when carrying relatively larger loads (Fig.3). Each factor by itself, head width (Estimate = 0.68, t_(808)_ = 1.86, *P* =0.06) and load area (Estimate = 0.11, t_(808)_ = 1.71, *P* =0.08), only marginally explained the number of antennae taps. When we analyzed the unladen ants, we found no relationship between the antennae taps per step and the head width (Estimate = -0.052, t_(798)_ = -0.33, *P* =0.73; Fig. 4), hence that factor by itself is not enough to explain the number of antennae taps.

**Figure 3.**
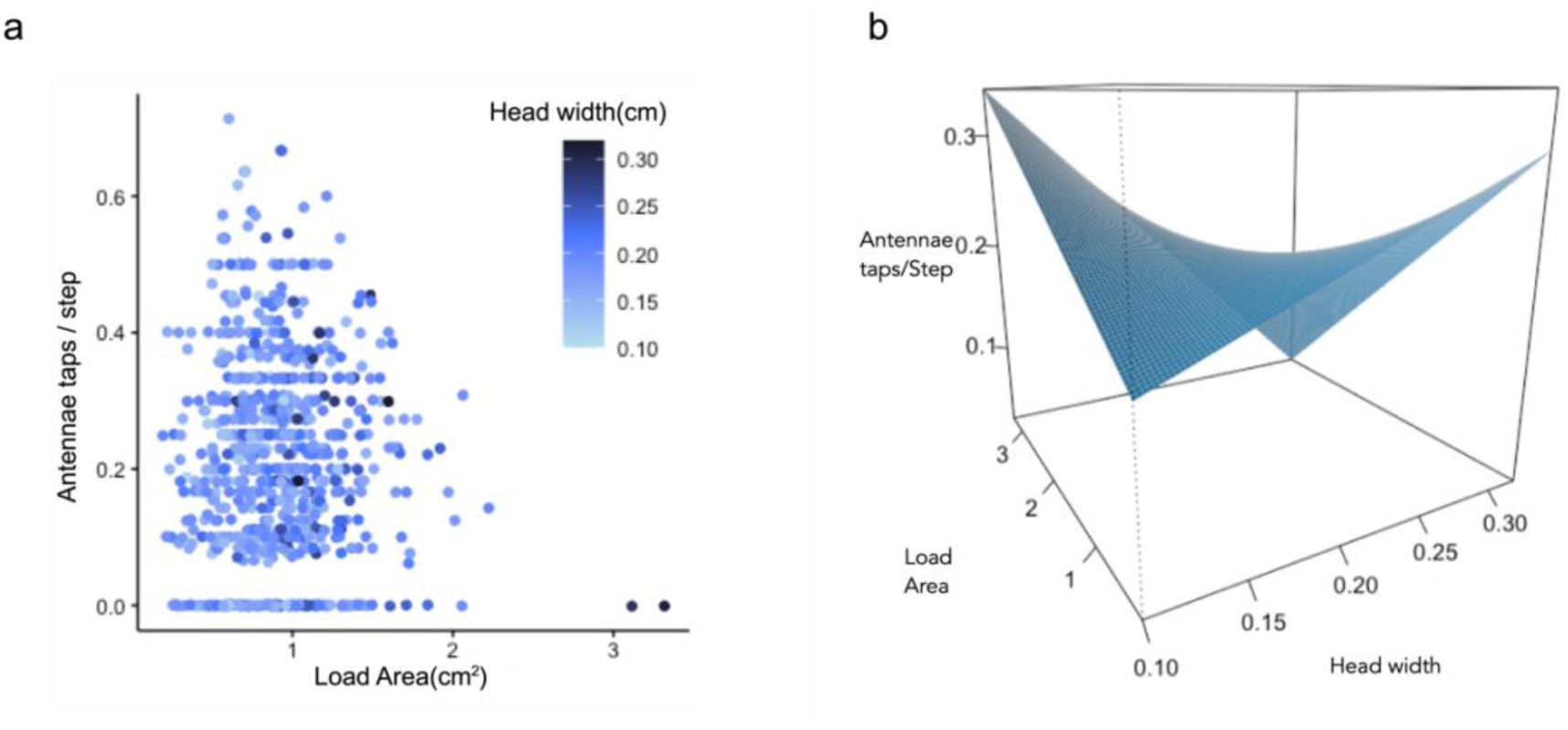
The number of antennae taps per step depends on the interaction between the load area and the worker’s size (head width). (**a**)The observed values were considered for an interaction, and we show (b) the plane of best fit according to the interaction of the two factors. As the interaction between the worker’s head width and the load size is significant, we can see that carrying an oversized load reduced the number of antennae taps for larger workers more strongly than for smaller workers.

**Figure 4.**
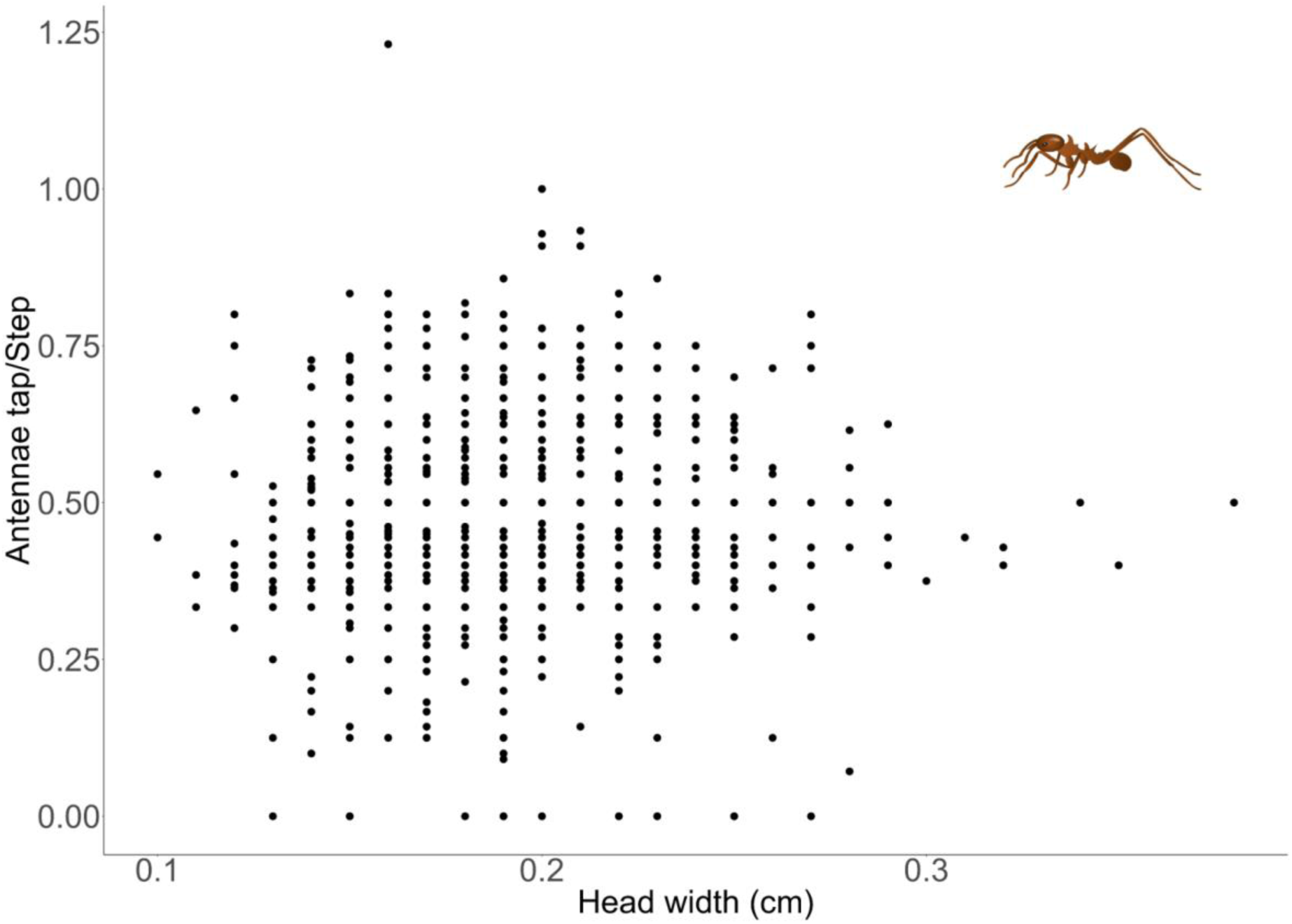
Number of antennae taps on the ground per step in non-laden leaf-cutter ants (*Atta colombica*) according to the worker’s head width. The worker’s size does not correlate with the number of antennae taps per step when ants are not carrying loads.

Conversely, the load shape did not affect the number of antennae taps (Estimate = - 0.03, t_(808)_ = -0.79, p=0.45), neither did the ant head width (Estimate = -0.11, t_(808)_ = -0.42, *P* =0.67), and there was no interaction between the factors (Estimate = 0.15, t_(808)_ = 0.66, *P* =0.50).

#### Antennae taps on nestmates

Laden ants contacted their nestmates from 0 to 4 times, and the number of contacts increased with the ant head width (Estimate=9.47, Z_(808)_=1.97, *P* =0.04), but not with the load area (Estimate=1.27, Z_(808)_=1.42, *P* =0.15; Fig. 5), neither with the interaction between both variables (Estimate=-5.29, Z_(808)_=-1.21, *P* =0.22).

**Figure 5.**
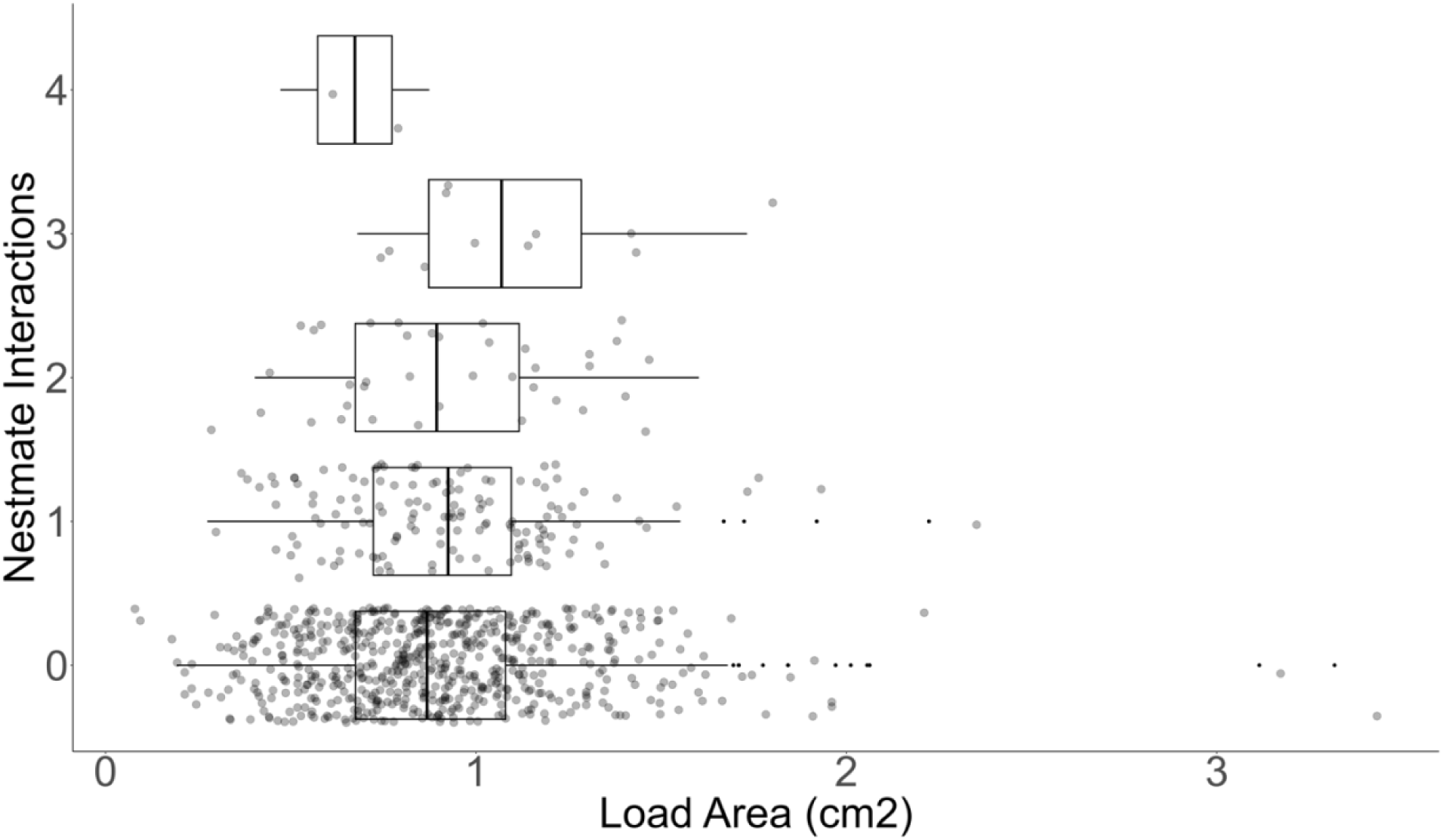
Number of nestmate interactions between laden and unladen leaf-cutter ants (*Atta colombica*) according to leaf fragment load area.

### Load reduction experiment

We recorded 44 ants before and after cutting the experimentally offered loads. We found that the number of antennae taps per step increased after experimentally reducing the load (X^2^=36.41, d.f.=1, *P* < 0.0001; Fig. 6a). Before the load reduction, the ants performed 0.26 (±0.12) antennae taps per step on average, compared to 0.39 (±0.14) after experimental load reduction. Workers also walked faster once the load was lighter: from 2.43 cm/s (± 1.15) to 2.96 cm/s (±1.32; X^2^=13.04, d.f.=1, *P* <0.001; Fig. S3a). Conversely, we recorded 33 ants for the control experiment, and we found that the number of antennae taps per step decreased from 0.39 (±0.16) to 0.35 (±0.16) antennae taps per step after touching the load with the dull scissors (X^2^=4.24, d.f.=1, *P* <0.05; Fig. 6b). The speed of the workers did not change after touching the load in control workers (before: 2.96 ±1.44 cm/s, after: 2.80 ±1.42 cm/s; X^2^=9.36, d.f.=1, *P* = 0.876; S3b).

**Figure 6.**
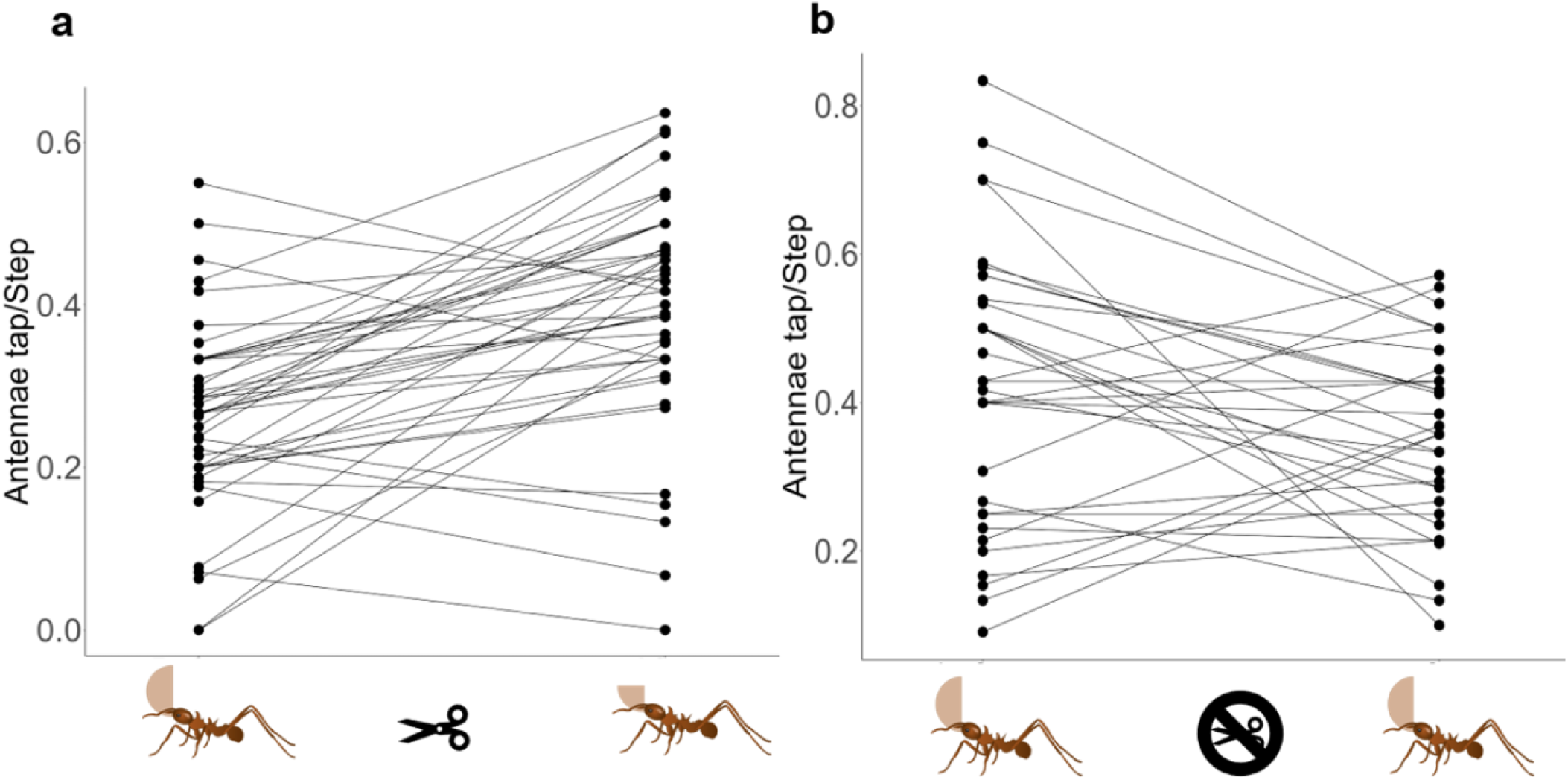
Number of antennae taps on the ground per step in the same individual of leaf-cutter ant (*Atta colombica*) **(a)** before and after reducing the load size by cutting the fragment in half, and **(b)** before and after touching the load with dull scissors as a control. Lines connect the paired values.

### Allometry of antennae length

The antennae length showed a negative allometry with head width (slope = 0.75, r= -0.55, d.f. = 98, *P* <0.0001; Fig. 7a), and thorax length (slope = 0.67, r= -0.67, d.f. = 98, *P* <0.0001; Fig. 7b). This means that larger ants (in thorax and head width) had proportionally shorter antennae than smaller ants. In contrast, other ant segments, such as the legs, grew isometrically to head width and thorax: first leg length changed isometrically with head width (slope = 1.00, r= 0.01, d.f. = 97, *P* = 0.88; Fig. S4a) and thorax length (slope = 0.89, r= -0.18, d.f. = 98, *P* = 0.07; Fig. S4b). The third leg also showed isometry with head width (slope = 1.05, r= 0.09, d.f. = 98, *P* = 0.35; Fig. S4c), and with thorax length (slope = 0.92, r= -0.12, d.f. = 98, *P* = 0.22; Fig. S4d).

**Figure 7.**
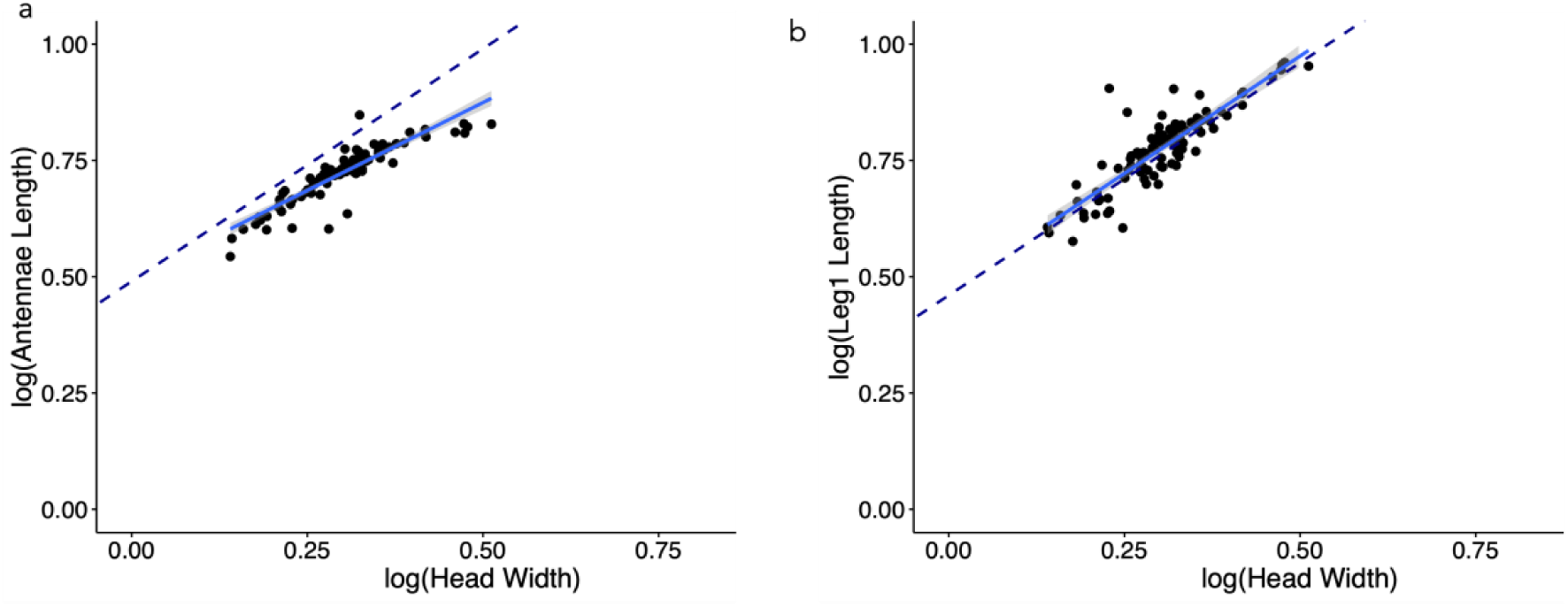
Relation between head width and (**a**) antennae length of leafcutter ant workers in log-log scale. The blue line depicts the slope of the association calculated using OLS, and the dotted line represents an isometric slope (slope=1), using the same intersection as the estimated equation. The antennae length showed negative allometry (a slope < 1). (**b**) A similar equation was calculated for the length of the first leg, which is isometric (slope = 1).

## DISCUSSION

We found evidence supporting the idea that carrying oversized loads negatively affects the behavior of tapping the ground with the antennae in leafcutter ants. First, we found that ant workers made fewer antennae taps per step when carrying a load than unladen ants. In *Atta vollenweideri,* workers carrying oversized loads uphill, horizontal, and downhill terrains do not balance the load by re-positioning in the mandibles, but instead by doing up and down movements of the head at the neck joint (Moll et al. 2010). In this study, the observations were done on flat surfaces, and workers still have to balance the fragment angle to keep the center of mass. Balancing the load by keeping a certain head position could be the reason for reducing the number of antennae taps while walking when carrying something in the mandibles. The head movements to balance the load may interfere with or limit the head or antennae movements for tapping the trail. Alternatively, unladen workers may perform more taps if searching for odors related to foraging decisions (Farji-Brener et al. 2010). However, our next finding challenges this explanation, because load size relative to worker size also affected the number of antennae taps.

The effect of the load size on the antennae taps per step depended on worker size. Larger workers did fewer antennae taps when carrying relatively larger loads than smaller ants also carrying oversized loads. Two reasons can explain this size-dependent effect. First, oversized loads in larger ants may be proportionally larger than for smaller ants, therefore limiting further the ability to perform antennae taps. However, our data shows that oversized loads (8 to 10 times larger than their ant head width) are found at any worker size (Fig. S2 and S5b). Alternatively, the greater effect on larger ants may be caused by the negative allometry between body size and antennae length, as larger ants have proportionally shorter antennae. This physical limitation, added to the head movements that large loads could impose to achieve balance (Moll et al., 2010), may incur in major sensorial impediment for laden larger ants towards the nest than for laden smaller ants. Additionally, in other species of leafcutters (*A. vollenweideri*), larger and smaller workers also have neuroanatomical differences in the brain antennal lobe, corresponding to differences in responses to trail pheromones (Kleineidam *et al*. 2007, Kuebler et al. 2010). Particularly, larger workers follow conspecific and heterospecific trails, whereas smaller workers preferentially follow conspecific trails. Further, contrary to smaller *A. vollenweideri* workers, larger workers have higher volume of glomeruli present in the first olfactory neuropil, the antennal lobe (Kuebler et al., 2010), which could be the case in the studied *A. colombica.* Beyond the potentially different sensorial abilities of different-sized workers, our data on the allometry of antennae length shows a physical limitation to tapping the ground in larger workers. Also, we cannot discard the possibility that larger ants hold their bodies at a greater distance from the ground or that having shorter antennae may require lowering the body more than a smaller ant would do to touch the ground with the antennae. The position of the body, combined with the negative allometry between worker size and antennae length, may result in larger ants having their antennae at a farther distance from the ground than smaller ants when carrying larger loads.

The first part of our study happened in a natural setting. Hence, leaf fragments may not come from the same leaf, individual plant, or plant species. Therefore, the load reduction experiment provides valuable evidence because it standardized the differences in loads. The experiment showed that the same individual would do more antennae taps if the load size allowed. After the experimental load reduction, most workers increased the antennae taps per step. Further, the increased tapping was related to an increase in the worker’s walking speed. Our results provided stronger evidence that load size affects the speed and the number of antennae taps per step and, therefore, the ability to smell the foraging trail. We also found differences in the number of antennae taps per step in our control, which deserves an explanation. Touching the load without cutting it caused a reduction in the number of antennae taps. This makes sense as touching the load interferes with the balance, and as we explained above, losing balance may force head movements that impede lowering the antennae. The effects that we see in the experiment are even more surprising because they would be the net result of reducing the antennae taps after the load was touched, and increasing them because they now carry a lighter weight. Previous reports demonstrated that the ants walking speed on the trails is decreased by higher load weight (Zollikofer, 1994; Rudolph & Loudon, 1986; Burd, 2000). Here, we propose that when carrying oversized loads, the limitation for smelling the trail (number of antennae taps) also causes workers to reduce the speed.

We found that the load shape did not affect the number of antennae taps per step, suggesting that other variables are more determinant in how the workers move their heads (e.g., balance in Moll et al. 2010). It has been reported for *A. vollenweideri* that workers were slower and consumed more energy when carrying long fragments of grass (elongated) than short fragments of grass of similar weights, presumably because workers carrying long fragments are running close to their stability limit (Moll et al., 2012). However, our study provides evidence that, in *A. colombica* ants, the shape of the leaf fragment does not consistently alter the ability of ant workers to tap the ground with the antennae. Since all our measurements come from workers walking on flat surfaces, we cannot discard that load shape may affect the number of antennae taps when ants walk in other terrains with different obstacles and slopes.

We also hypothesized that tapping the nestmates to acquire information could compensate for any limitation related to tapping the ground with the antennae. Although the number of interactions between laden and unladen ants was not associated with load area, it was a relatively common behavior in our results. Hence, workers may be often using their nestmates to acquire information to orientate in the trail. Besides pheromones, leaf-cutter ants could use visual cues to stay on the foraging trail, because they can perceive some movements (Vilela, Jaffé & Howse, 1987), and workers could also use information from magnetic fields to orientate (Banks & Srygley, 2003; Riveros & Srygley, 2008; Riveros et al., 2014). Additionally, in other ant species (*Lasius niger*), individuals deposit fewer pheromones during the day than at night, indicating that light may help them visually orientate (Jones et al., 2019). Another factor that should be considered is the biochemistry of trail pheromones. They are known to be a mixture of volatile, semi-volatile, and non-volatile compounds (Tumlinson et al., 1972; Jaffé et al. 2011). Hence, the air may be a useful agent in the chemical reception of pheromones in the trail, making it less necessary to put the antennae on the ground when they are carrying something, and successfully reduce the effort or save energy. However, studies regarding the need to touch the surface to smell the trail pheromone vary on the conclusion depending on the study species. For instance, *Camponotus* ants need to touch the surface to smell the trail pheromones (Draft et al. 2018), but *Monomorium* ants have short-lasting pheromones they can detect in the air (Robinson et al. 2008). *Atta cephalotes* cannot orient back to the nest without trail pheromones (Wetterer et al. 1992), but the biochemical composition and detection distances of trail pheromones are unknown in *A. colombica*.

More generally, our study was aimed to answer the question of why workers would carry loads smaller than what they can lift. If they carried the maximum weight they could, they colony would be getting a larger amount of food per foraging trip. Our results show a potential limitation related to sensorial impediments. Another explanation is that workers may leave room for the extra weight they get on the trip, for instance, if the load becomes wet or if nestmate hitchhikers climb on the leaf (Linksvayer et al., 2002). Other selective forces may also explain their ability to lift heavy loads, even when this is not used for foraging. For instance, lifting heavy obstacles from the foraging trail may require a strong force over a short distance, where the impediment to smell the foraging trail would not have greater consequences for the ant worker. Workers often abandon these obstacles near the trail and unladen when navigating back to the foraging trail. Moreover, orientation, when removing very large obstacles, seems to be based on communication with nestmates (Bochynek et al., 2019; Alma et al., 2020) instead of relying on trail pheromones. Larger ants of *A. colombica* participate more frequently in removing obstacles from the trail (Howard, 2001). Hence, large workers may lift heavy objects or manipulate fragments of larger areas without the need to constantly smell the trail.

Our observations of ants carrying loads in nature evidenced that smaller and larger ants with small loads may have fewer difficulties tracking the pheromones back to the nest. Our load reduction experiment showed that one ant individual can adjust their movements according to the properties of the load. Although it is known that “truck drivers” (i.e., workers with oversized loads) are not common in trails with high traffic because of the use of space (Pereyra & Farji-Brener, 2020), another potential reason for choosing medium-sized loads in nature is a lower cost of detecting the chemical trail. The individual decisions of ants are crucial for the colony’s success, because a single action can impact or influence the whole foraging operation of the superorganism. These results show how workers assess the risks or costs of slowly moving an oversized load compared to moving a smaller load at a faster speed. How this variation in the antennae taps per step affects the worker’s ability to navigate or orientate back to the nest is a question that deserves further studies.

Understanding the sensorial mechanisms and cognitive processes of animals have helped illuminate our understanding of foraging behavior and ecology. Here, we provide an example where a limitation on the ability to sense the world imposed by the foraging task itself, may explain foraging decisions that do not fit optimal foraging theory expectations. Similar mechanical limitations in the sensory system are probably faced by other social and solitary organisms while foraging, and documenting these limitations will help us further our understanding of foraging decisions.

## Data availability

Data is available in FigShare DOI 10.25573/data.25762197

