## Supplementary Materials for "Carrying oversized loads may create pheromone “blind spots” in leafcutter ants"

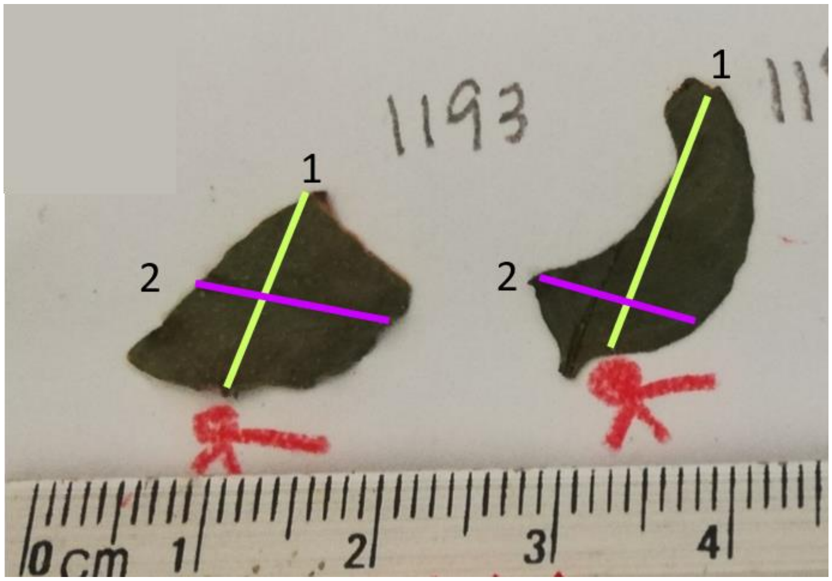


**Figure S1.** Load shape was calculated by marking the leaf fragment by the place the ant was holding it with the mandibles (red mark) based on the videos and drawing a line to the furthest edge (load length, line 1 in yellow). Load width (line 2 in purple) was a perpendicular line to length encompassing the widest part of the leaf.

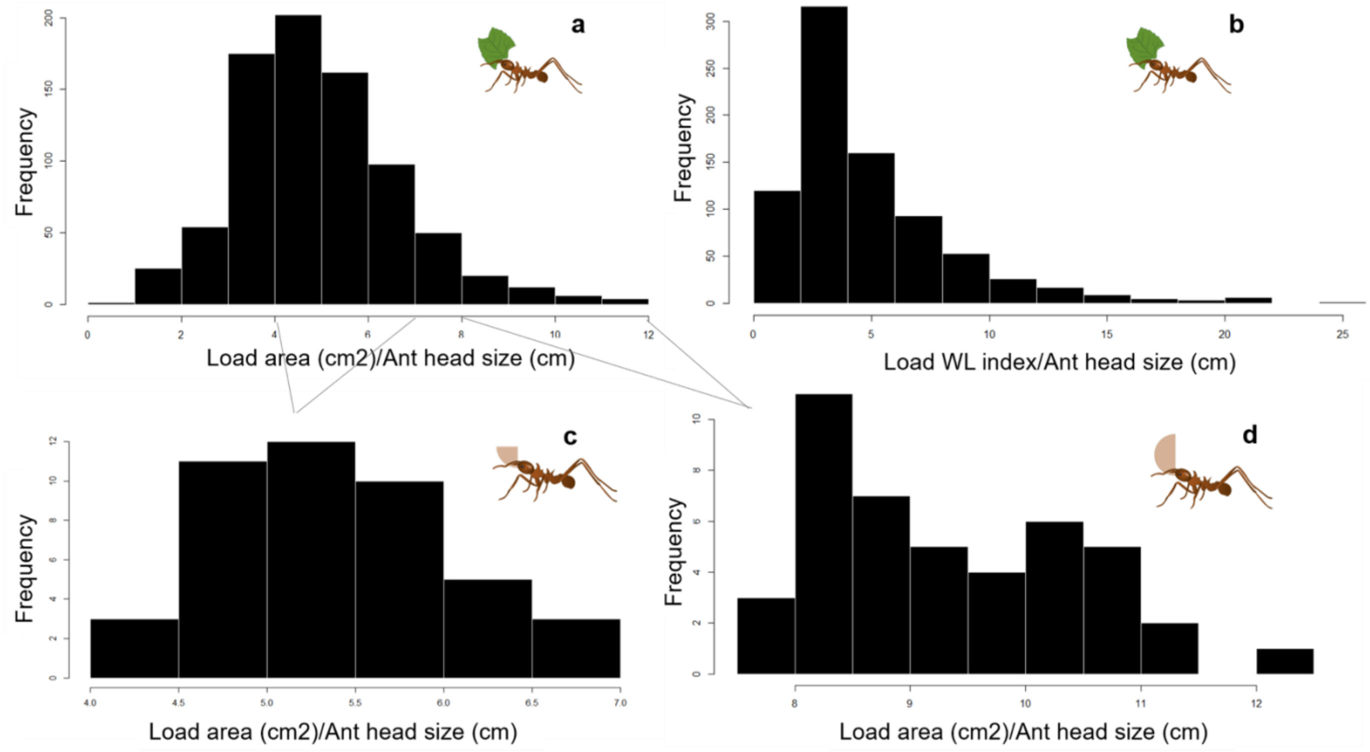


**Figure S2.** Number of laden leaf-cutter ants (*Atta colombica*) carrying loads of different **(a)** area and **(b)** shape (width to length index) in natural conditions, corrected by ant head width (See the section of "Orientation to the nest according to the load" in Methods). In the load reduction experiment, we are showing how ants fit in the distribution of the naturally occurring loads **(c)** after reducing the experimental load and **(d)** before reducing its size, considering head width (See the section "Plasticity in the load transport to the nest" in Methods). Note that workers who picked up the experimentally offered load carry an oversized load compared to what is found in nature ("truck drivers"). After experimentally reducing the load, these ants carried medium-size load in nature.

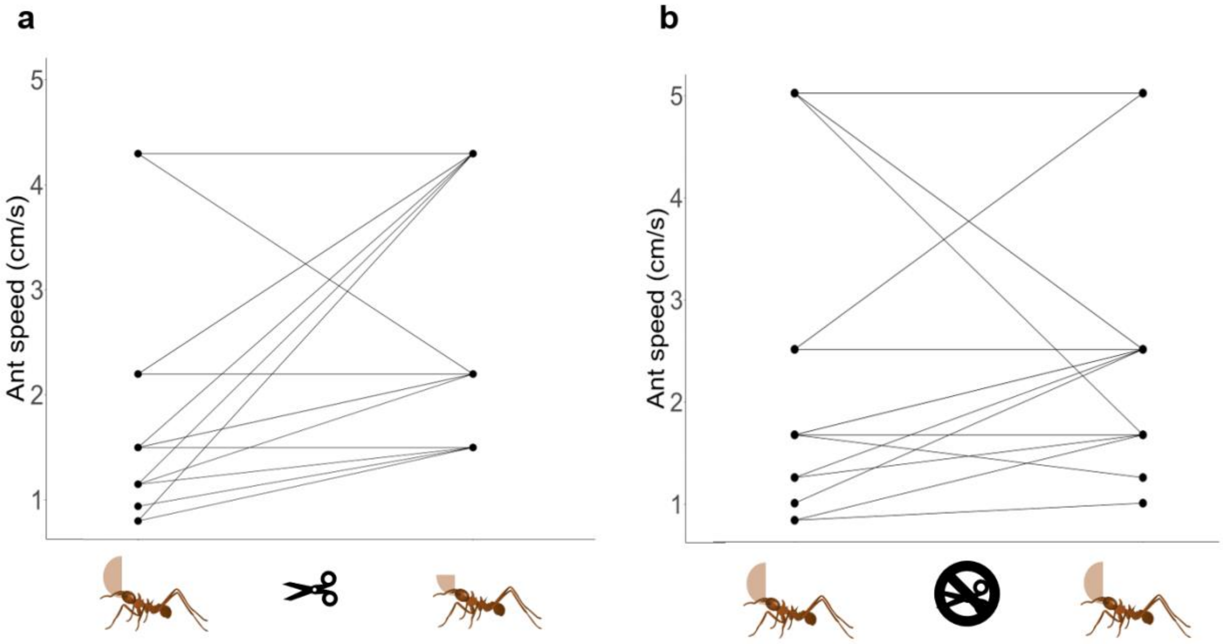


**Figure S3.** Speed of leaf-cutter ants (*Atta colombica*) **(a)** before and after reducing the load size by cutting the fragment in half, and **(b)** before and after touching the load with the dull scissors as a control treatment. Lines connect the paired values.

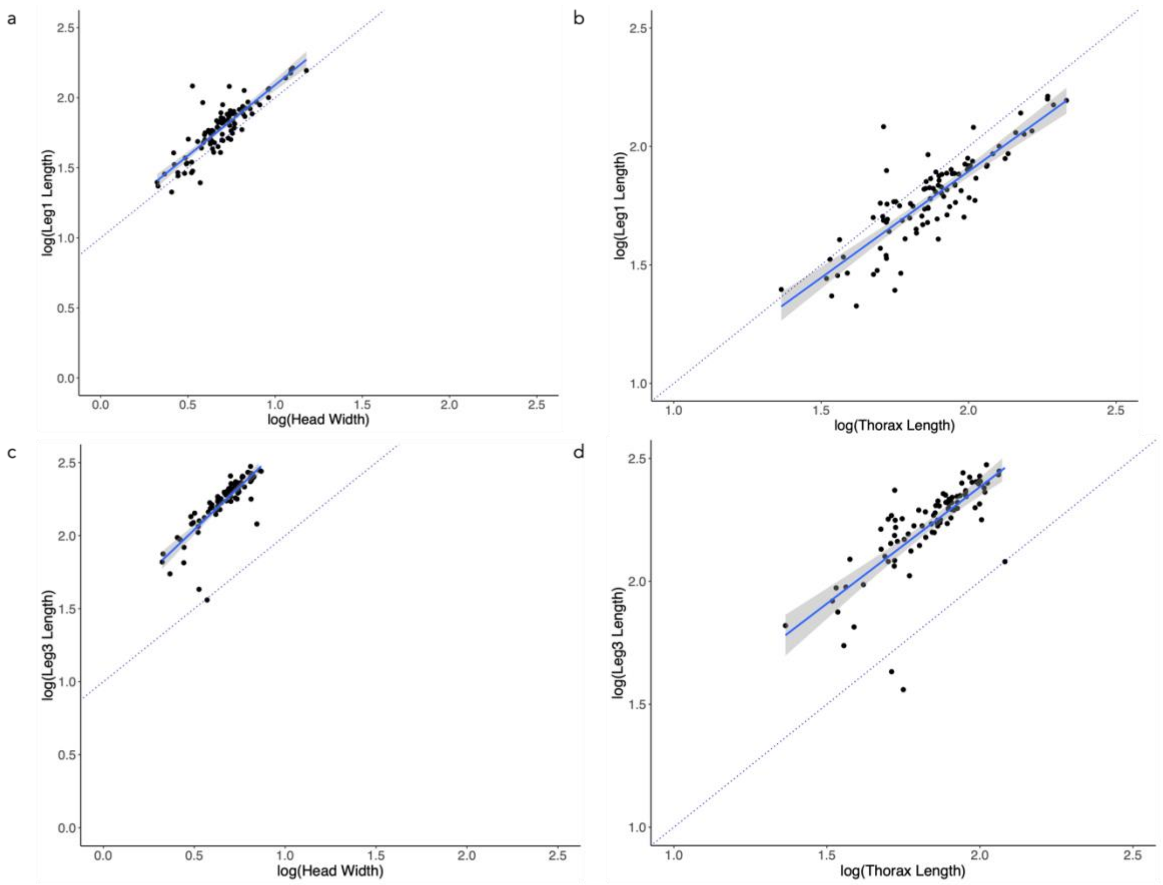


**Figure S4.** Relation between leafcutter ant worker first leg length with (**a**) head width or (**b**) thorax length, and third leg length with (**c**)head width and (**d**) thorax length. The blue line depicts the slope of the association calculated using OLS, and the dotted line represents an isometric slope (slope=1).

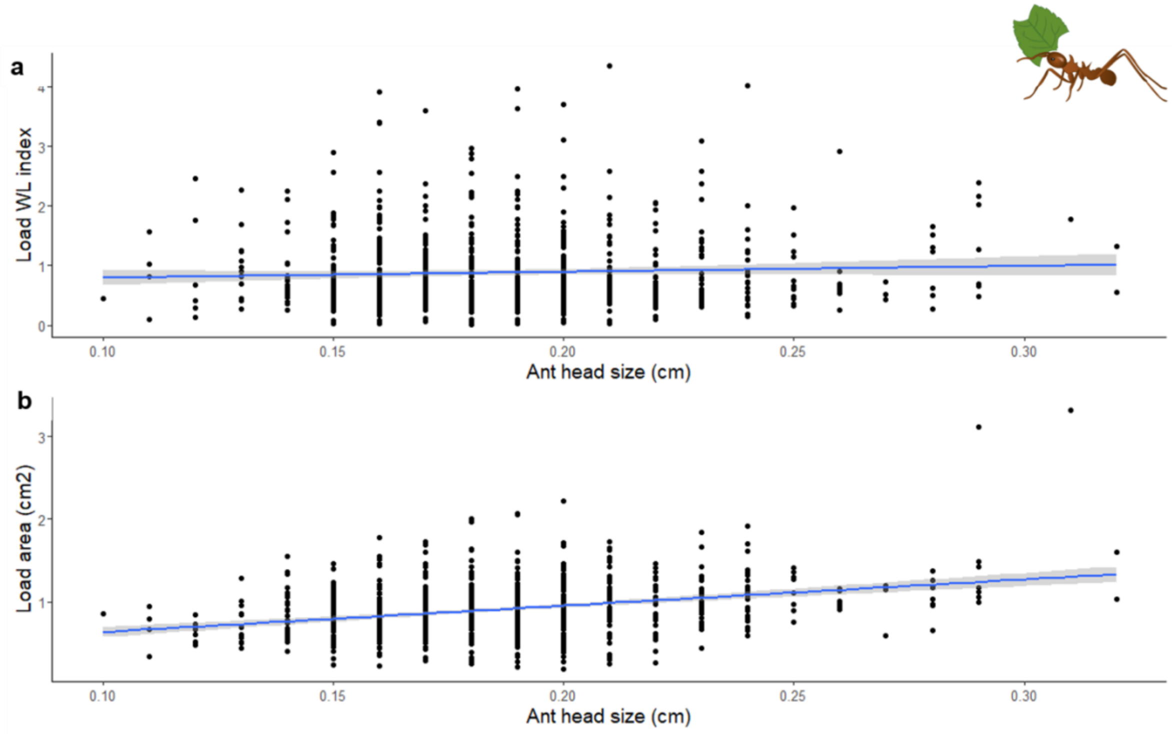


**Figure S5.** Leaf fragments properties carried by leafcutter ants (*Atta colombica*) according to their head width. To understand whether larger ants were carrying larger loads, we performed two generalized linear models (1) between the ant head width and the load area and (2) between the ant head width and the load shape, using the WL index. We found that **(a)** there was no relationship between the ant head width and the shape of the load (Estimate=0.99, $t_{(808)}$=1.45, *P* =0.15; Fig. S3a), and **(b)** larger ants did transport loads of larger area (Estimate=3.18, $t_{(808)}$=9.26, *P* <0.0001).
